# Pathogen and endophyte assemblages co-vary with beech bark disease progression, tree decline, and regional climate

**DOI:** 10.1101/2021.02.24.432696

**Authors:** Eric W. Morrison, Matt T. Kasson, Jeremy J. Heath, Jeff R. Garnas

## Abstract

Plant-pathogen interactions are often considered in a pairwise manner with minimal consideration of the impacts of the broader endophytic community on disease progression and/or outcomes for disease agents and hosts. Community interactions may be especially relevant in the context of disease complexes (i.e, interacting or functionally redundant causal agents) and decline diseases (where saprobes and weak pathogens synergize the effects of primary infections and hasten host mortality). Here we describe the bark endophyte communities associated with a widespread decline disease of American beech, beech bark disease (BBD), caused by an invasive scale insect (*Cryptococcus fagisuga*) and two fungal pathogens, *Neonectria faginata* and *N. ditissima*. We show that the two primary fungal disease agents co-occur more broadly than previously understood (35.5% of infected trees), including within the same 1-cm diameter phloem samples. The two species appear to have contrasting associations with climate and stages of tree decline, wherein *N. faginata* was associated with warmer and *N. ditissima* with cooler temperatures. *Neonectria ditissima* showed a positive association with tree crown dieback – no such association was observed for *N. faginata*. Further, we identify fungal endophytes that may modulate disease progression as entomopathogens, mycoparasites, saprotrophs and/or additional pathogens, including *Clonostachys rosea* and *Fusarium babinda*. These fungi may alter the trajectory of disease via feedbacks with the primary disease agents or by altering symptom expression or rates of tree decline across the range of BBD.

## 1. Introduction

Plant-microbe or plant-insect interactions are often considered in a pairwise manner with minimal consideration of the impacts of the broader community on the nature and outcomes of herbivory or pathogen attack. In some systems, tri- or even multipartite interactions have been shown to be important, particularly where fungus-insect or fungus-insect-mite symbioses are involved (Wingfield et al., 2010, 2016). Rarely, however, are broader communities considered, despite the fact that pathogens and insects attacking forest trees are embedded in variable and/or, in the case of non-native species, novel communities. In the most aggressive and well-studied examples (e.g., Chestnut blight, Dutch elm disease, or the more recent Emerald ash borer), disease agent aggressiveness and ensuing host mortality may be sufficiently rapid such that co-occurrence with other organisms is of minimal relevance to system dynamics, though colonization with certain endophytic microbes can influence host susceptibility (Feau and Hamelin, 2017) or pathogen aggressiveness (Kolp et al., 2020) in subtle or complex ways. Often, however, interactions with host trees are embedded in a diverse community that varies both spatially and temporally, and diseases “complexes” (diseases with multiple causal agents that act in concert to produce symptoms and mediate host decline), are increasingly recognized as important (Desprez-Loustau et al., 2016). The role of the community in determining host fate is extremely difficult to ascertain and could proceed via multiple, potentially interacting mechanisms (Table 1).

**Table 1.** Summary of ecological guilds in addition to primary pests and pathogens that are potentially involved in tree decline associated with multi-species disease complexes. These include both insects and fungi and may have negative (-), positive (+) or no effect (0) on hosts.

| Functional role (guild) | Effect on host | Description |
| --- | --- | --- |
| Other pathogens | - | Direct effects on host, could synergize or antagonize primary insect/pathogen impacts |
| Decay fungi | -/0 | Increase susceptibility to snap, interrupt vascular transport, reduce resource quality or availability for primary insect/pathogen (could be positive of host) |
| Other insects | -/0 | Competition for resources, tree defense induction, predation (including intra-guild), vectors (for other microbes, mites, nematodes, etc.) |
| Mycoparasites | + | Reduce growth, survival/longevity, or spore production of 1°/2° pathogens |
| Entomopathogens | +/0 | Reduce survival, longevity of 1°/2° insects |
| Non-pathogenic microbes | +/-/0 | Competition for resources, tree defense induction, viral reservoirs? |
| Endophytes | +/-/0 | Roles variable and largely unknown, potential defensive symbionts, latent pathogens, early-colonizing saprotrophs, etc. |

Beech bark disease (BBD) in North America is a canker disease of American beech (*Fagus grandifolia*) caused by an invasive scale insect, *Cryptococcus fagisuga* Lind., and one of two presumptively native fungal pathogens, *Neonectria faginata* and *N. ditissima*. BBD is referred to as a disease complex because it involves both insects and fungi with at least some degree of ecological redundancy with respect to disease etiology and symptom development. The invasive felted beech scale (*C. fagisuga*) is recognized as the primary initiating agent of BBD symptoms, but only inasmuch as it facilitates infection of beech trees by the fungal BBD pathogens *Neonectria faginata* and *N. ditissima* (Ehrlich, 1934). Both *N. faginata* and *N. ditissima* cause similar cankering infections and bole defects resulting in host callus tissue formation (Cotter and Blanchard, 1991). In addition to the primary disease agents, at least two fungal mycoparasites are known from the BBD system – *Clonostachys rosea* (the anamorph and preferred name of *Bionectria ochroleuca*; Houston et al., 1987; Schroers et al., 1999; Rossman et al., 2013; Stauder et al., 2020a) and *Nematogonum ferrugineum* (Houston, 1983). Other fungi are regularly isolated from infected trees (i.e., *Fusarium babinda*; Stauder et al., 2020a) though their roles are less clear. Further, saprotrophic fungi are likely involved in late stages of disease wherein host stem tissue is weakened to the point of mechanical failure (“beech snap”, Houston, 1994), though the presence or importance of wood rot fungi on disease progression is unknown.

In addition to questions of the role of the broader community in disease dynamics, the relative frequency and ecological importance of the two primary (*Neonectria*) pathogens has long been the subject of study and debate. While apparently ecologically similar within the BBD system, the life histories of these fungi differ in important ways. For example, *Neonectria ditissima* is a generalist that infects many diverse tree hosts including species of birch, maple, walnut, mountain ash, and holly, among others (Castlebury et al., 2006; Stauder et al., 2020a). The species is also an important pathogen of apple (Gomez-Cortecero et al., 2016). The diversity and abundance of alternative hosts could plausibly influence ecological and evolutionary dynamics (Houston, 1994; Kasson and Livingston, 2009), though this question has not been adequately studied to date. In contrast, *N. faginata* has never been observed outside of the BBD complex in North America (Castlebury et al., 2006) despite recent surveys focused on uncovering possible cryptic native reservoirs for this pathogen (Stauder et al., 2020a).

These fungi exhibit spatiotemporal trends with respect to the timing of site-level infestation with the felted scale. The current range of beech bark disease is defined by the range of the felted beech scale, which, unlike many forest pests, has spread slowly (∼13 km per year; Morin et al., 2007) from the site of initial introduction in Halifax, Nova Scotia in 1890. This progression, together with a handful of long-distance dispersal events (i.e., to North Carolina, West Virginia, and Michigan) has resulted in a gradient of duration of infestation ranging from very recent (∼9 years in Wisconsin) to more than eight decades (86 years in Maine) (Houston, 1994; Cale et al., 2017). Surveys of *Neonectria* species distribution have generally found *N. ditissima* to be more prevalent in the killing front of the disease (i.e., 10-20 years post scale insect arrival; Cale et al., 2017 and references therein). *Neonectria faginata* appears to dominate aftermath forests to the degree that researchers have suggested near replacement of *N. ditissima* with *N. faginata* as early as seven years after pathogen attack becomes apparent (Houston, 1994; Cale et al., 2017). However, *N. ditissima* can maintain a presence in stands dominated by *N. faginata*, including within the same tree (Kasson and Livingston, 2009). Persistence of *N. ditissima* in areas that are previously BBD-affected may be attributed to reinfection from reservoirs of this fungus in non-beech hardwood tree hosts (Kasson and Livingston, 2009), and/or the development of secondary killing fronts when climatic conditions allow beech scale to colonize areas where it was previously inhibited (e.g., release from killing winter temperatures by warm periods; Kasson and Livingston, 2012). These species – while morphologically indistinguishable in the field – can be separated using culture morphology and spore size measurements, though the latter process is tedious and is dependent on the presence of sexually produced ascospores. Further, spore size comparisons can only detect co-infection if either both species are simultaneously producing perithecia (sexual spore structures) or multiple isolates are collected, cultured, and induced to mate and produce sexual structures (Cotter and Blanchard, 1981; Stauder et al., 2020b). Partly because of these challenges, it is yet unclear whether trends in species dominance with infection duration are consistent, whether *N. ditissima* plays an important and/or predictable role in aftermath forests, and how the prevalence and distribution of each species reflects climate, disease stage, and other tree host and environmental conditions.

The objectives of this study are twofold. First, we examine patterns of occurrence (and co-occurrence) of *N. faginata* and *N. ditissima* across the current range of BBD and use joint species distribution modeling to evaluate hypothesized biotic and abiotic drivers of the prevalence and relative dominance of these species. Second, we characterize the bark mycobiome of American beech using Illumina-based metabarcoding on bark samples collected from across the range of BBD to ask how disease-associated communities vary geographical and to assess the fidelity and potential role of key species within the BBD system, whether as direct or indirect drivers or as indicators of disease state. Based on previously observed patterns with respect to disease dynamics across the range of BBD, we evaluated the relative contribution of hypothesized drivers of pathogen and associated community distribution, including duration of regional infection with BBD, disease severity (i.e., tree condition as well as beech scale and *Neonectria* perithecia density), and climate.

## 2. Methods

### 2.1 Site description and sample collection

Bark disks including phloem tissue were collected from American beech (*Fagus grandifolia*) from ten sites across the range of BBD. Sites ranged from northern Maine, western North Carolina, and eastern Wisconsin, representing latitudinal and longitudinal transects across the current range of BBD with a range of infection duration (Table 2). Sampling was performed from December 2017 through January 2019. At each site American beech trees were surveyed for levels of *Neonectria* perithecia density, beech scale density, crown dieback, and amount of cankering. Scale insect and *Neonectria* fruiting density were scored on a 0-5 ordinal scale (Houston et al., 2005; Garnas et al., 2011a) and tree condition on a 0-4 scale. Distinct cankering types were pooled and measured as roughly corresponding to 20% bins by bole coverage. Trees were sampled along 100 x 5 m transects in a random direction from starting point up to a maximum of 50 trees or 400 meters. All trees were measured along the transect to facilitate estimation of tree density and size distribution, etc. Where possible, stratified random sampling was performed for bark plug collection so as to obtain an unbiased sample across a range of tree conditions. Stratified sampling levels were tree size (three levels 1^st^, 2^nd^/3^rd^, and 4^th^ quartile of diameter at breast height [DBH]), four levels of *Neonectria* perithecia density (0, 1, 2-3, 4-5), and two levels of beech scale density (0-1 v. 2-5), yielding 24 possible stratification levels. It was not possible to collect all combinations at all sites, but most sites had representative trees in most categories. In particular, in one site (Wisconsin) there were no visible *Neonectria* perithecia and we instead stratified within DBH and the available levels of beech scale density (2-3, 4-5) with four replicates per stratum (n = 24). Bark plugs were collected using a flame-sterilized 1-cm diameter hollow leather punch and stored on ice in sterile 24-well plates. Multiple plugs were taken from a random subset of trees with number of plugs ranging from 1-6. Samples were stored on ice and then frozen within 48 hours of sampling and stored at -20°C until processed for DNA extraction.

**Table 2.** Site locations and duration of BBD infection in terms of years since beech scale first observed. Beech scale observations based on Cale et al., 2017.

| State | Latitude | Longitude | Number of trees with ≥ 1000 sequences | Duration of infection (years) |
| --- | --- | --- | --- | --- |
| Maine | 46.6585 | -68.6913 | 10 | 68 |
| Maine | 44.8307 | -68.5996 | 7 | 68 |
| New Hampshire | 43.1340 | -70.9510 | 13 | 59 |
| New York | 44.4924 | -74.0295 | 6 | 43 |
| New York | 43.0841 | -74.4406 | 11 | 58 |
| Pennsylvania | 41.2144 | -75.3834 | 8 | 42 |
| Michigan | 45.3179 | -84.6723 | 11 | 12 |
| Wisconsin | 44.9284 | -87.1891 | 21 | 9 |
| West Virginia | 38.6074 | -79.8443 | 10 | 19 |
| North Carolina | 35.3203 | -82.8439 | 5 | 20 |

We used the PRISM dataset (PRISM Climate Group, 2020) to calculate climate variables for each site. We first determined the start and end dates of the growing season in each year based on empirical values describing heat accumulation for American beech leaf out and leaf drop (Richardson et al., 2006), wherein bud break for a given site and year was estimate as the date when a site had accumulated 100 cumulative GDD_4_ (base 4°C) from January 1. Leaf drop was defined by 500 cumulative chilling degree days (below 20°C) from August 15. These calculations resulted in leaf out estimates ranging from March 15 to May 12 and leaf fall estimates from October 20 to November 8 along the natural climate gradient among sites. We considered nongrowing season climate because fungi are likely to grow in periods where minimum temperatures are non-limiting, and growth during periods of tree host dormancy may be important for fungal establishment, growth, and/or aggressiveness. We then summed GDD_4_, daily precipitation, and freeze-thaw frequency (the number of days that temperatures crossed 0°C) for both the growing season and nongrowing season.

### 2.2 DNA extraction, PCR, and sequencing

Phloem plugs were prepared for DNA extraction by removing surface debris, *Neonectria* perithecia, and the periderm layer using a sterile scalpel. Plugs were then washed under a steady stream of 1 ml sterile 1x PBS pH 7.2 buffer. Phloem plugs were freeze-dried for 24 hours then crushed and homogenized, and a subsample of phloem (mean 56 mg ±1 mg standard error) was subjected to bead bashing. DNA was extracted from ground phloem samples using a QIAGEN DNeasy Plant Mini Kit following the factory protocol. Extracted DNA was then purified using a Zymo OneStep PCR Inhibitor Removal Kit following the factory protocol and diluted 1:10 in PCR grade water.

The ITS2 region was amplified in duplicate PCR reactions with the primers 5.8S-Fun and ITS4-Fun using Phusion High Fidelity polymerase, a 58°C annealing temperature, and 30 PCR cycles (Taylor et al., 2016). Primers included the Illumina TruSeq adapters and sample identification tags were added to amplicons via a second round PCR at the University of New Hampshire Hubbard Center for Genome Studies. A dual-unique indexing strategy was used such that each sample had a matching pair of indices on forward and reverse reads in order to reduce potential for sample misassignment due to index switching or index bleed. We included PCR negatives (one per PCR plate) and DNA extraction negatives (one per kit) in the sequencing run.

### 2.3 Bioinformatic analyses

Amplicon sequence variant (ASV) calling was performed using the DADA2 (Callahan et al., 2016) protocol v1.8 for ITS sequences in R 3.6.2 (DADA2 version 1.14.0), with an additional step to extract the ITS2 region from 5.8S-Fun–ITS4-Fun amplicon sequences using *itsxpress* (Rivers et al., 2018). The core DADA2 algorithm was run with option ‘pool = TRU’ to allow greater sensitivity for ASVs that were rare in a single sample but more abundant across the entire dataset, and putative chimeras were removed using DADA2::*removeBimeraDenovo*. Sequences from trees with multiple plugs were pooled at the tree level before ASV calling. Taxonomy was assigned to ASV representative sequences by comparison to the UNITE dynamically clustered database (release date 04/02/2020; Nilsson et al., 2019) using the DADA2::*assignTaxonomy* algorithm.

We used the LULU post-processing algorithm to group putatively erroneous ASVs with their parent ASVs based on sequence similarity and co-occurrence patterns (Frøslev et al., 2017), which has been shown to improve reconstruction of biological taxa for fungi relative to ASV denoising alone (Pauvert et al., 2019). We note that this is not a clustering algorithm but rather uses minimum sequence similarity as the first of three criteria for culling of likely error variants. After comparing a range of minimum sequence similarity thresholds (84%, 90%, 93%, 95%) we selected a 93% threshold, which maximized the identification and removal of likely erroneous ASVs while minimizing taxonomic reassignment.

One of the goals of this study was to determine the distribution of *N. faginata* and *N. ditissima* across the range of BBD. In order to increase sensitivity for the detection, after calling ASVs we subsequently mapped the original quality filtered reads to representative sequences for ASVs identified as *N. faginata* and *N. ditissima* using the ‘vsearch usearch_global’ algorithm (Rognes et al., 2016). This approach increases detection sensitivity by including reads that may have otherwise been discarded or misassigned during ASV calling steps (Edgar 2013; Pauvert et al., 2019). Samples were scored as containing *N. faginata* or *N. ditissima* if the species were discovered in a sample using either the ASV calling or mapping-based approaches. Singleton ASVs were removed from the dataset and samples with less than 1000 sequences after singleton filtering were also excluded.

### 2.4 Statistical analyses

We first performed pairwise correlations of site characteristics data to examine relationships between disease severity and climate variables using the R package Hmisc (Harrell et al., 2019). These were either means of disease severity variables collected from random transects, or 10-year mean climate variables for each site extracted from the PRISM database. We first tested for normality using a Shapiro-Wilk test (Shapiro and Wilk 1965). We report Spearman rank correlation (ρ) where one or both variables was non-normal (including all ordinal variables) – for other variables we report Pearson correlation (r).

To test for deviations from random patterns of co-occurrence of the two *Neonectria* species we used a probabilistic model (Veech 2013; Griffith et al., 2016). In brief, all possible permutations of species occurrence were determined based on the number of samples and observed species frequencies. A probability distribution was then calculated wherein the probability of a given co-occurrence frequency was equal to the number of permutations with that co-occurrence frequency as a proportion of the total possible permutations. Significant deviations from random were then assessed by comparing the observed co-occurrence to the probability distribution with *P* equal to the sum of probabilities for co-occurrence in less than (*P_lt_*) or greater than (*P_gt_*) the observed number of samples. We excluded samples from the Wisconsin site, which at nine years post-beech scale colonization had a considerable number of uninfected trees that would have skewed the analysis toward detecting aggregation.

To examine the effects of environmental covariates on *N. faginata* and *N. ditissima* occurrence, we applied spatially explicit joint species distribution modeling implemented in the R HMSC package (Ovaskainen et al., 2016; Ovaskainen et al., 2017; Tikhonov et al., 2017). We used the *N. faginata* and *N. ditissima* presence-absence matrix as the dependent variables and employed a probit link function. Pairwise site geographic distance was included as a random effect to control for spatial correlations between predictors and *Neonectria* occurrence. Tree was included as nested random effect (within site); log(sequence count) was included as a fixed effect to account for sampling depth effects on species detection probability (Ovaskainen et al., 2017). Non-growing season climate variables were stronger predictors of *Neonectria* occurrence in exploratory analyses, and we therefore only considered nongrowing season variables in subsequent models. The primary model included climate variables and disease severity variables in order to determine the best predictors of *N. faginata* and *N. ditissima* occurrence. Independent variables were standardized to their respective means and standard deviations to obtain comparable slope estimates. We report Tjur’s *R*^2^, a coefficient of determination for logistic regression, (Tjur, 2009; Ovaskainen et al., 2017) along with slope coefficients. To minimize issues with multicollinearity, precipitation was excluded from the model due to strong correlations with beech scale density (ρ = -0.88, *P* < 0.001) and DBH (ρ = -0.73, *P* = 0.02, Supplemental Table S1).

We used PERMANOVA (*vegan::adonis* function, Oksanen et al., 2019) to determine whether *Neonectria* species occurrences were associated with differences in composition of the remaining community. We used the presence-absence of each ASV with greater than 10% frequency as predictors of community composition, by first randomly subsampling the dataset to 1000 sequences per sample, iteratively removing each predictor ASV from the dependent variable matrix, and then performing PERMANOVA on transformed sequence counts (log_10_+1) with the removed ASV as a categorical predictor. We tested for a relationship between ASV frequency and its strength as a predictor of community composition by regressing PERMANOVA *R*^2^ against ASV sample incidence.

We next used indicator species analysis to explore whether certain species were associated with *Neonectria* species occurrence, *Neonectria* perithecia density, beech scale density, crown dieback, or cankering using the *indicspecies::multipatt* function (De Caceres and Legendre, 2009). We chose these variables because in combination they describe various stages of tree decline and aspects of disease and are amenable to transformation to categorical predictors (and therefore appropriate for ISA analysis). We used the sample-ASV matrix, randomly selecting 1000 sequences per sample for this analysis. We then identified ASVs that were indicators of at least two measures of disease severity. The goal of filtering out ASVs that were indicators of only one disease severity measure was to increase interpretability and reduce misclassification of ecological roles due to spurious correlations. We then performed functional classifications of ASVs using the FungalTraits database (Põlme et al., 2021). We noted primary and secondary lifestyles, and where additional functional potential was noted, such as endophytic capacity, we recorded this as tertiary lifestyle. We also noted “wood saprotroph” as a separate category, but collapsed other saprotrophic categories (i.e., “unspecified saprotroph,” “litter saprotroph,” “soil saprotroph”) into a single “saprotroph” designation.

### 2.5 Data accessibility

Raw sequence data and associated sample metadata are archived at the NCBI SRA under BioProject accession PRJNA701888.

## 3. Results

### 3.1 Disease severity and site characteristics

We performed a pairwise cross-correlation analysis using site-level means of disease severity variables and climactic variables (Table S1). As expected, there were strong correlations between climate variables and latitude. Specifically, latitude was negatively correlated with heat accumulation (GDD_4_) in both the growing (r = -0.64, *P* = 0.047) and nongrowing season (r = -0.95, *P* < 0.001) and with precipitation in the growing season (Spearman’s ρ = -0.96, *P* < 0.001). Latitude was also negatively correlated with elevation (r = -0.73, *P* = 0.016) with more southerly sites tending to be at higher elevation. Longitude was not significantly correlated with the climate variables tested but was strongly positively correlated with duration of BBD infection (r = 0.97, *P* < 0.001) and cankering (ρ = 0.83, *P* = 0.003) and negatively with elevation (r = -0.69, *P* = 0.03). Various climate parameters correlated with disease severity. Nongrowing season precipitation was negatively correlated with both scale insect density (ρ = -0.88, *P* = 0.001) and DBH (ρ = -0.73, *P* = 0.02), while growing season precipitation was negatively correlated with scale insect density (ρ = -0.81, *P* = 0.005). We found no other significant correlations between climate and disease severity indicators. There were, however, correlations among some of the disease severity indicators we measured. Duration of infection correlated positively with cankering (ρ = 0.81, *P* = 0.004), while scale insect density and DBH were also positively correlated (ρ = 0.65, *P* = 0.043). No other correlations with disease severity indicators were found. We note, however, that the youngest site (Wisconsin, infested in 2010) had high mean wax density but no visible *Neonectria* perithecia at the time of sampling. We performed pairwise partial-correlation analysis using nongrowing season precipitation, DBH, and beech scale while controlling for the effect of the third of these variables in each pairwise combination. Partial correlation analysis showed that only nongrowing season precipitation and beech scale retained a significant correlation after controlling for the effect of DBH (ρ_precip,scale.DBH_ = -0.78, *P* = 0.013), whereas the other relationships were no longer significant (ρ_scale,DBH.precip_ = 0.12, *P* = 0.97; ρ_precip,DBH.scale_ = -0.45, *P* = 0.22).

### 3.2 Sequencing results

After sequence processing and ASV denoising the number of sequences per sample (tree) ranged from <100 to 331,624 (median 21,315) across 117 samples. The LULU post-processing algorithm resulted in the identification of 149 ASVs (13% of 1132 original ASVs) that were better interpreted as variants of existing “parent” ASV’s (either due to minor sequence variation or sequencing error; Supplementary Table S2). Over 90% of the ASVs culled in this way shared full taxonomic identity with their respective parent ASVs. For example, the dominant *N. faginata* and *N. ditissima* ASVs each subsumed 8 and 3 “children” variants respectively with no taxonomic reassignment. Fourteen ASVs were taxonomically reassigned, but in all cases this involved reassignment to a higher taxonomic rank (i.e., species designations were removed but the genus [13 ASVs] or family [1 ASV] was retained). After LULU post-processing and removing samples with less than 1000 sequences 796 ASVs remained across 102 samples retained, with the number of ASVs per sample ranging from 12 to 172 (median 60). Rarefying (to 1000 sequences per sample) led to a further drop in ASV retention; ASVs richness ranged from 7 to 67 per sample (median 25). The mean ASV richness per site ranged from 13.2 ± 4.38 s.d. to 46.8 ± 16.4 s.d. and total site richness ranged from 66 to 412 ASVs. Further exploration of the relationships between disease severity, climate, and broad trends in community composition and diversity are possible using this dataset and are currently underway. We focus here on the primary fungal disease agents and description of species that may play a role in disease progression (Table 1).

### 3.3 Neonectria species distribution and its drivers

*Neonectria faginata* was present in all ten sites and *N. ditissima* was found in all but the southernmost site in North Carolina (Fig. 1). Importantly, the two species often co-occurred both at the site level and within the same tree, including within the same 1-cm phloem disc. Overall, *N. faginata* was present in 135 of 170 phloem plugs (79.4%) and 71 of 102 trees (69.6%), while *N. ditissima* was present in 46 of 170 plugs (27.1%) and 32 of 102 trees (31.4%). The two species co-occurred in 38 of 170 plugs (22.4%) and 27 of 102 trees (26.5%) including 35.5% of the 76 trees infected with at least one species. Both species were absent in 27 of 170 plugs (15.9%) and 26 of 102 trees (25.5%). At least one of the two species was detected in all 109 plugs where perithecia were present on the plug periderm surface prior to processing for DNA extraction (64.1% of plugs). *Neonectria faginata* was detected in 107 (98.2%) of the plugs with fruiting structures present while *N. ditissima* was detected in 31 plugs (29.4%). The species co-occurred in 29 of 109 plugs (26.6%). Sixty plugs had no perithecia present and one plug of the 170 total had degraded periderm such that it was not possible to record perithecia presence-absence. Of the 60 plugs with no perithecia at least one species was detected in 33 plugs (55%), wherein *N. faginata* was detected in 27 (45%) and *N. ditissima* in 15 (25%). The two species co-occurred in nine plugs without perithecia (15%), and were both absent in 27 plugs (45%).

**Figure 1.**
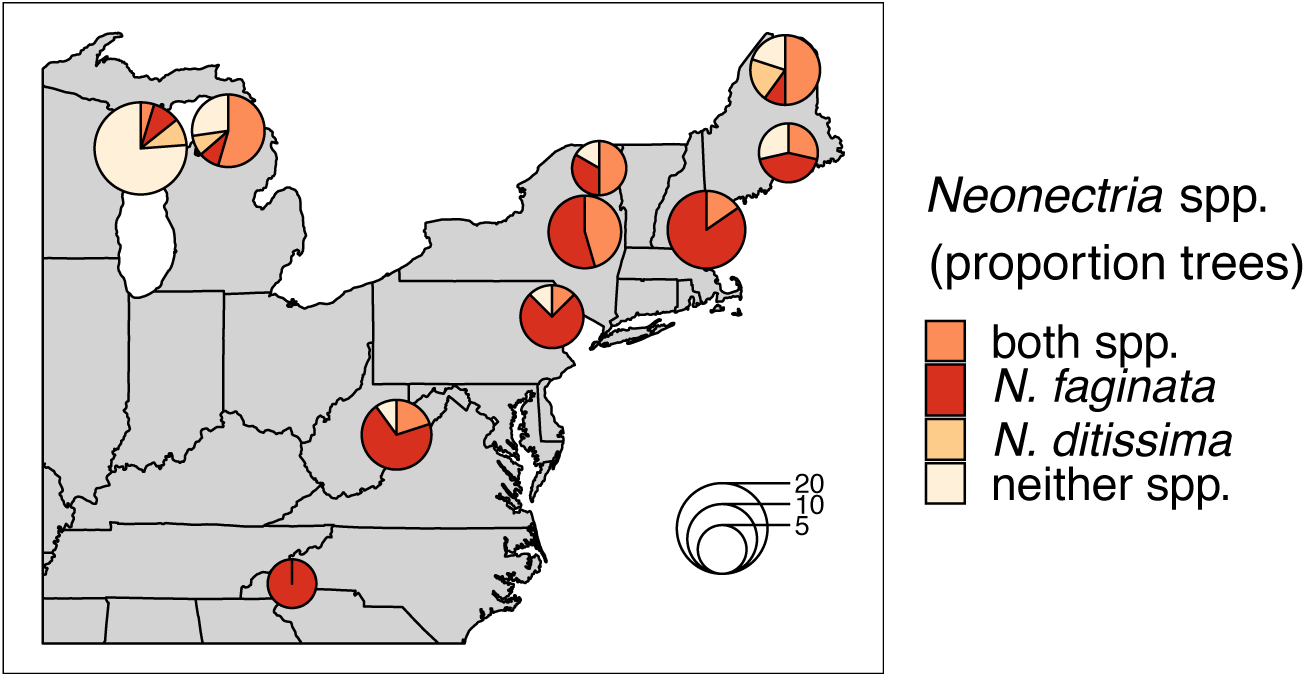
*Neonectria* species occurrence across ten sites. Proportions in pie charts indicate the number of trees in a site where a species was detected using metabarcoding. The number of trees per site is indicated by pie chart diameter.

*Neonectria ditissima* was only found in isolation at the tree level (i.e., without *N. faginata*) at the three northernmost sites we sampled (MI, WI, and northern ME). In the remaining seven of the ten sites, *N. ditissima* was only detected in trees where *N. faginata* was also present. In terms of species prevalence, *N. faginata* occurred in a greater number of trees than *N. ditissima* in all but the three northernmost sites (90% mean occurrence versus 25% mean occurrence for *N. faginata* and *N. ditissima*, respectively, in the seven more southerly sites). The two species each occurred in isolation in two trees in Wisconsin and co-occurred in one tree, and in Michigan each species occurred in isolation in one tree and co-occurred in six trees. *Neonectria ditissima* was more prevalent in our northern Maine site (70% versus 60% of trees for *N. ditissima* and *N. faginata*, respectively, including 50% of trees where the species co-occurred). In sites within the aftermath zone (i.e., excluding the Wisconsin site within “advancing front” of the disease), *N. faginata* was detected in 84% of all trees across our sites, whereas *N. ditissima* was detected in 36% of trees. Given these observed occurrence frequencies, co-occurrence did not differ from a random distribution (32% observed vs. 30% expected; *P_gt_* = 0.24 *sensu* Veech [2013]).

We next examined correlations between *Neonectria* species incidence across sites (i.e., presence-absence at the tree level) and indices of disease severity and climate using spatially explicit joint species distribution modeling (HMSC, Ovaskainen et al., 2017). We first tested for effects of growing season versus nongrowing season climate parameters and found that climate during the nongrowing season was overall a stronger predictor of patterns of *Neonectria* occurrence (Supplemental Fig. S1). Specifically, heat accumulation (GDD_4_) during the nongrowing season was significantly associated with incidence of both species, while growing season GDD_4_ was generally a poor predictor of incidence. Nongrowing season freeze-thaw was significantly associated with *N. faginata* (posterior support *P* = 0.99) but not *N. ditissima* incidence (*P* = 0.86). In the growing season neither of these variables was correlated with *Neonectria* species occurrence.

After controlling for sampling effort and spatial structure, our models explained 54% and 37% of variance in *N. faginata* and *N. ditissima* incidence, respectively, based on Tjur *R*^2^ (Tjur, 2009). The full model indicated that *N. faginata* had a significant, positive relationship with nongrowing season heat accumulation (*R*^2 =^ 0.19), duration of infection (*R*^2^ = 0.14) and DBH (*R*^2 =^ 0.04), and a negative association with beech scale density (*R*^2 =^ 0.06) (Fig. 2). Heat accumulation was the strongest predictor of *N. faginata* incidence (*R*^2^ = 0.19). *Neonectria ditissima* incidence was positively associated with DBH (*R*^2^ = 0.07), freeze-thaw cycle frequency (*R*^2^ = 0.07), and crown dieback (*R*^2 =^ 0.04), but negatively associated with nongrowing season heat accumulation (*R*^2^ = 0.07) and beech scale density (*R*^2^ = 0.05). Results were unchanged when nongrowing season precipitation was included in the model, except there was no significant correlation between *N. faginata* and beech scale or infection duration (Fig. S2). Neither species showed a significant correlation with nongrowing season precipitation. The HMSC approach also allows examination of residual correlation between dependent variables (i.e., *N. faginata* and *N. ditissima* occurrence) after accounting for the effect of independent predictors. We found no residual correlation between the two species after accounting for the effects of disease severity and climate, and also found no correlation between the species using a reduced model that only controlled for sampling effort (i.e., sequence count) and spatial structure. This result suggests that distribution of the two species is not strongly structured by inter-species interactions (e.g., competition or facilitation).

**Figure 2.**
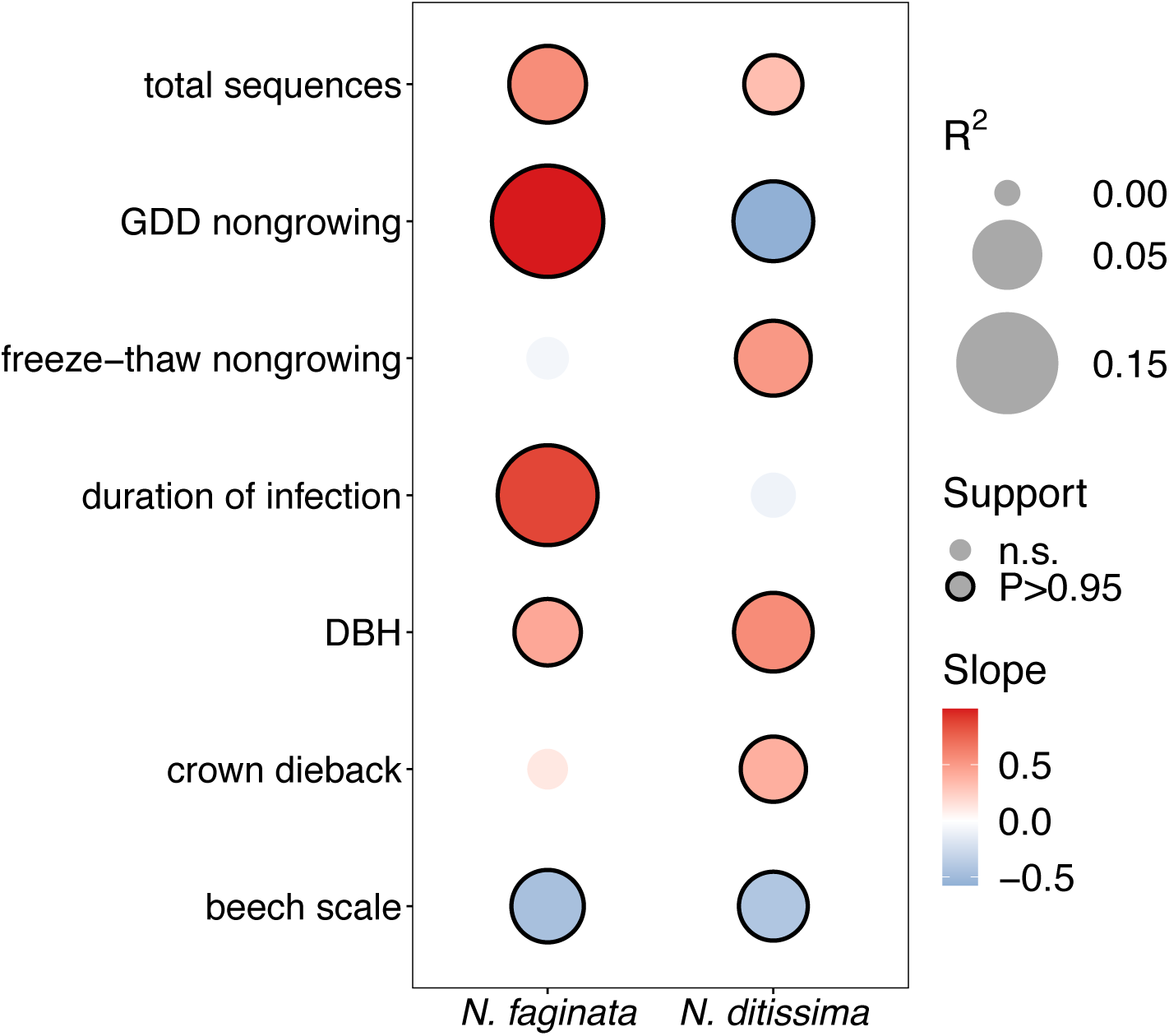
Effects of disease severity and climate on *Neonectria* species occurrence within trees. Points with solid outlines represent significant effects at posterior probability *P* > 0.95 (analogous to a p-value of *P* < 0.05). Box fill indicates the strength and direction of the relationship with blue indicating a negative slope, and red indicating a positive slope. Point size indicates magnitude of *R*^2^ with overall fits for *N. faginata* and *N. ditissima* of Tjur *R*^2^ = 0.54 and 0.37, respectively.

In light of the significant positive association of *N. ditissima* with crown dieback indicated in HMSC modeling, we visually explored patterns of species incidence across levels of crown dieback and cankering (Fig. 3). *Neonectria ditissima* occurrence increased across crown dieback classes (crown dieback level 0 = 21%, 1 = 24%, 2 = 38%, 3 = 58%), whereas *N. faginata* occurrence was relatively stable or modestly declined (crown dieback level 0 = 79%, 1 = 71%, 2 = 62%, 3 = 67%). The increase in *N. ditissima* with increasing crown dieback resulted in an increase in co-infection by the two species at the tree level. *Neonectria ditissima* occurred in isolation in 2-8% of trees in each dieback class, whereas trees co-infected with both *Neonectria* species increased across dieback classes (crown dieback level 0 = 16%, 1 = 21%, 2 = 31%, 3 = 50%). When discrete cankers were absent both species were at relatively low occurrence (38% and 26% for *N. faginata* and *N. ditissima*, respectively). However, *N. faginata* more than doubled in trees with cankers compared to no cankers (85% *versus* 38%, respectively; χ^2^ = 21.6, *P <* 0.001), and *N. ditissima* doubled in the highest cankering level compared to levels 0-2 (50% *versus* 25%, respectively; χ^2^ = 5.1, *P =* 0.02) resulting in elevated co-infection at the highest cankering category (46% of trees compared to 17-23% in remaining categories).

**Figure 3.**
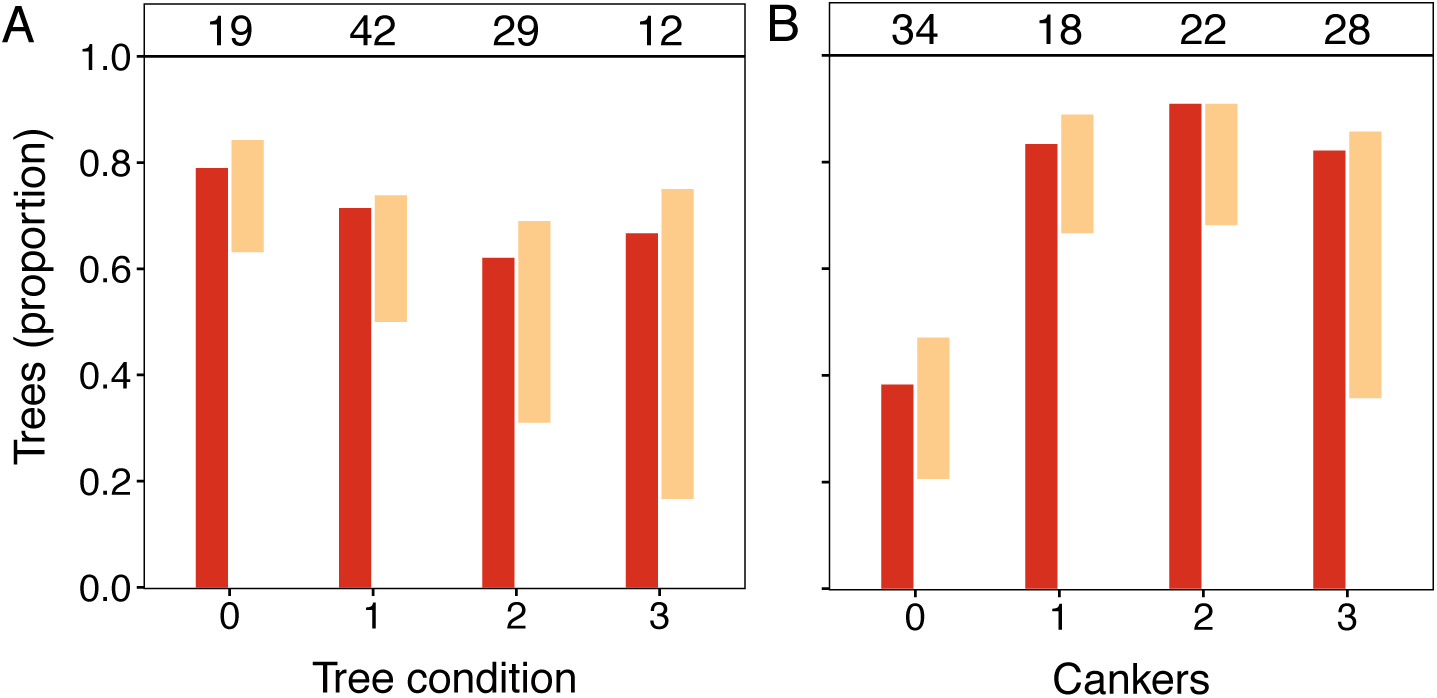
Occurrence of *N. faginata* and *N. ditissima* across levels of crown dieback (A) and cankering (B). The proportion of trees in which each species was detected is indicated for *N. faginata* (dark red) and *N. ditissima* (light red). The proportion of co-occurrence is indicated by overlap of the bars and the remaining white space indicates the proportion of trees where neither species was detected. The number of trees per grouping (n) is indicated above respective bars.

In this work we were also interested in whether *N. faginata* or *N. ditissima* was structuring or responding to different fungal endophyte communities. Of the 62 most commonly detected ASVs (minimum 10% incidence across all samples) *N. faginata* was the fifth best predictor of mycobiome composition (PERMANOVA *R*^2^ = 0.076, *P* = 0.001) while *N. ditissima* was the 42^nd^ best predictor (*R*^2^ = 0.023, *P* = 0.007; Supplemental Fig. S3A). We tested for a relationship between PERMANOVA *R*^2^ and sample incidence to examine the possibility that species presence-absence effects on community composition were primarily driven by the frequency of ASV occurrence. There was a weak but significant relationship between incidence and PERMANOVA *R*^2^ (*R*^2^ = 0.165, *P* = 0.001; Supplemental Fig. S3B) with the top five most predictive ASVs occurring in between 13% and 61% of samples.

### 3.3 Ecological roles of fungi in the BBD system

We used a combination of Indicator Species Analysis (ISA; de Caceres and Legendre 2009) and literature-based functional classification (Põlme et al., 2021) to discern degrees of statistical association and potential ecological roles for fungal species that appear as beech bark endophytes across the range of disease and tree decline levels sampled. Overall, 38 ASVs were identified as indicator species of at least two disease categories. All but two of these ASVs were indicators of different levels of crown dieback or cankering (Table 3). Nine of the 38 were indicators of the absence of crown dieback and either low levels of cankering or the absence of scale insect (see Table 3, “Healthy beech” indicators). Of these nine, only three were taxonomically identified to the genus level, with six other ASVs identified to order, phylum or kingdom. Another five ASVs were indicators of both low levels of cankering and absence of scale insect (Table 3, “Minor cankering, scale absent”), with two being taxonomically identified to the genus level.

**Table 3.**
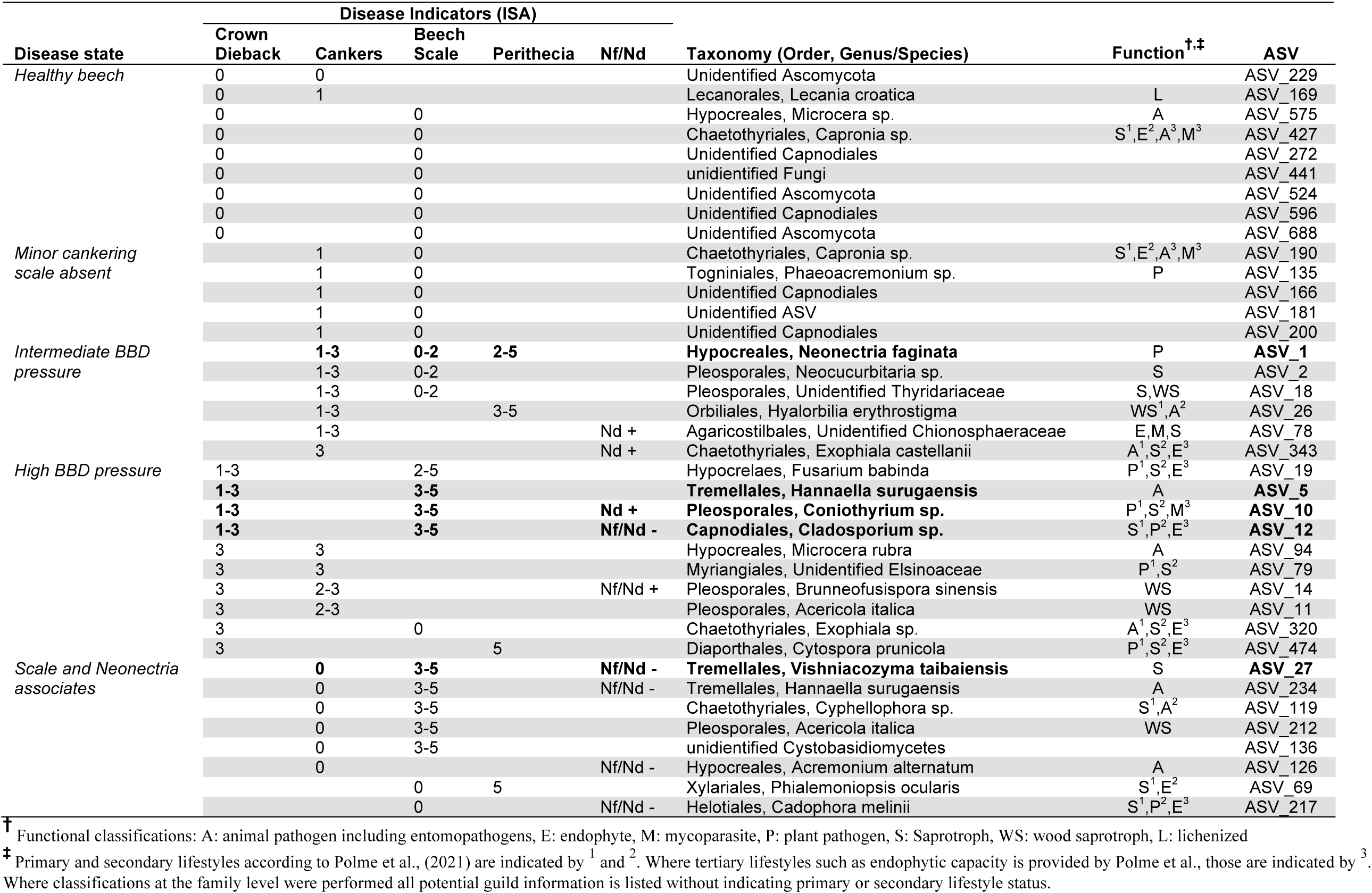
ASVs associated with at least two indicators of disease according to ISA (columns “Disease indicators (ISA)”). Taxonomy is listed as order, species epithet where available, or most informative taxonomic level. Those five ASVs that were strongest predictors of community composition according to PERMANOVA are bolded. Functional classifications are provided (*sensu* Põlme et al., 2021).

In total, 16 ASVs were associated with intermediate to high levels of either cankering or crown dieback. Six ASVs were indicators of intermediate to high levels of cankering, low scale density, and/or presence of *Neonectria* species (Table 3, “Intermediate BBD pressure”). These included *N. faginata*, as well as ASVs annotated as animal pathogens/entomopathogens, mycoparasites, plant pathogens, and saprotrophs or wood saprotrophs, the latter two functional groupings accounting for five of the six ASVs. Ten ASVs were indicators of intermediate to high levels of crown dieback (Table 3, “High BBD pressure”). Four of these ten were also indicators of high scale density, and another four were indicators of high levels of cankering. Eight of the ten ASVs associated with high levels of crown dieback were annotated as either saprotrophs or wood saprotrophs, and five were annotated as plant pathogens. All five of the ASVs assigned a plant pathogen function mapped to multiple functional groups, with saprotrophic lifestyle assigned as primary or secondary functions. Three ASVs associated with high levels of crown dieback were also among the top five predictors of community composition (Supplemental Fig. S3), and were annotated to entomopathogen (animal pathogen), plant pathogen-saprotroph-mycoparasite, or saprotroph-plant pathogen-endophyte functions, respectively. We note that *N. ditissima* was also an indicator of high levels of crown dieback (crown dieback 2-3) but was not formally included in this analysis due to lack of association with additional disease indicators.

Eight ASVs were associated with presence or absence of the primary disease agents (Table 3, “Scale and *Neonectria* associates”), including ASVs associated with high beech scale density and *Neonectria* absence (two ASVs), high density of both beech scale and *Neonectria* perithecia (one ASV), or absence of both beech scale and *Neonectria* (one ASV). Six of the eight were also associated with absence of cankering. Five of these eight ASVs were annotated to saprotroph or wood saprotroph functions, three as entomopathogens/animal pathogen, and two as endophytes, including three ASVs with multiple functional mappings.

We also specifically explored the distributions of *Clonostachys rosea, Nematogonum ferrugineum*, and *Fusarium babinda* given their previously described roles as BBD associates as either mycoparasites of *Neonectria* (*C. rosea* and *N. ferrugineum*; Barnett and Lilly 1962; Houston, 1983; Stauder et al., 2020a) or potential entomopathogens or secondary beech pathogens (Stauder et al., 2020a). Two ASVs were annotated as *C. rosea* and together they occurred in four of our ten sites ranging from 10% to 27% of trees in respective sites (Fig. 4A). One of these ASVs was also an indicator of the highest level of *Neonectria* perithecia. One ASV of putative importance as a mycoparasite of the BBD fungi, *N. ferrugineum,* only occurred at two sites at low frequency (9 and 20% of trees Fig. 4B) and was not a statistical indicator of any disease categories. Another ASV (ASV 19) was identified as *Fusarium babinda*, which has been previously identified in association with BBD and is a suspected beech scale associate (Stauder et al., 2020a). This ASV was associated with high wax density and high crown dieback (Table 3) and occurred in all ten of our sites (Fig. 4C).

**Figure 4.**
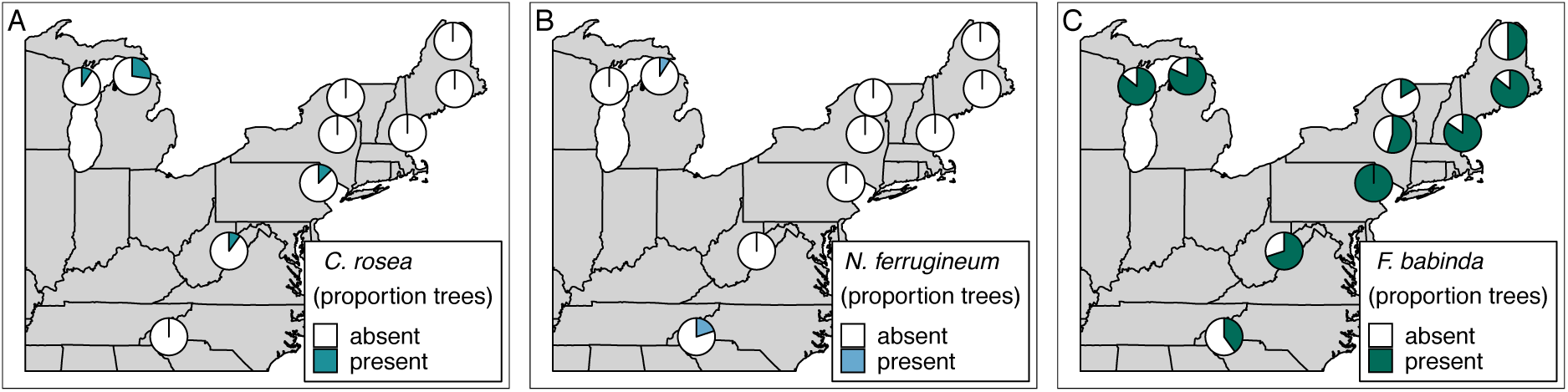
Occurrence of ASVs 16 and 21, *Clonostachys rosea* (A), ASV 807, *Nematogonum ferrugineum* (B), and ASV 19, *Fusarium babinda* (C), across ten sites. Proportions in pie charts indicate the number of trees in a site where the species was detected using metabarcoding.

## 4. Discussion

In the present study we explored the distribution of *N. faginata* and *N. ditissima*, the primary pathogens involved in BBD, in relation to disease severity and climate characteristics in ten sites across the range of BBD. We further explored patterns of association with other fungal species in the beech bark endophytic community in relation to tree disease state, and used these relationships to highlight and hypothesize ecological roles of potential relevance to BBD severity and progression. We show that *N. faginata* and *N. ditissima* have divergent correlations with climate and may be associated with different stages of tree decline. Further, we show that fungal species occurring in association with the primary BBD agents (i.e., the host, *F. grandifolia*; the scale insect initiating agent, *C. fagisuga*, and the fungal pathogens, *N. faginata* and *N. ditissima*) may contribute to disease outcomes in complex and interacting ways in this system. We suspect that these types of feedbacks are not unique to BBD dynamics but rather are likely to generalize to other complex tree diseases.

### 4.1 Correlations among disease agents and climate

We found that precipitation in both growing and nongrowing seasons was negatively correlated with scale insect density. Nongrowing season precipitation correlated negatively with DBH. The former of these correlations is consistent with a hypothesized causal relationship whereby precipitation reduce scale insect population density by washing colonies off of tree boles (Houston and Valentine, 1988; Dukes et al., 2009; Garnas et al., 2011b; Kasson and Livingston, 2012). The latter relationship between precipitation and DBH is likely driven by multicollinearity between infection duration (and thus disease severity indicators), geographic distribution, and climate. Sites with older infections tend to be dominated by small diameter trees and also occur in lower precipitation sites in our dataset. That said, cankering (which constitutes evidence of past, non-lethal infection) was the only index of disease status that was related statistically (positively in this case) with duration of infection. Our dataset included sites ranging from nine to 68 years from the arrival of beech scale based on county-level data (Cale et al., 2017), but only one site was in the “advancing front” stage of infection as typically defined (i.e., ≤10 years post scale insect arrival; Houston et al., 2005). At least one further site, and possibly up to three sites, occurred in the “killing front” stage of infection (i.e., 5-10 years after advancing front conditions, or 15-20 years after scale insect arrival; Houston et al., 2005), whereas the remaining sites were sampled in an aftermath forest infection stage. Interactions between disease agents are variable in aftermath forests despite density dependent growth within populations of insects or fungi (Garnas et al., 2011a). Our sampling design, which was weighted towards aftermath forest stands, may have obscured the typically observed patterns across disease stages in beech scale and fungal pathogen abundance. However, there were signals of duration of infection-driven patterns in our dataset. For example, our advancing front site had high mean wax density but no visible *Neonectria* perithecia production, as is typical of advancing front stage forests (Shigo, 1972; Garnas et al., 2013). Further, scale insect density and DBH were positively correlated, largely because sites with younger infections – and thus higher beech scale density associated with earlier disease stages – also tended to have larger trees. Relationships between scale insect densities within the aftermath forest alone are considerably weaker (Garnas et al., 2011b).

### 4.2 Neonectria species distribution and its drivers

Both *Neonectria* species were widely distributed geographically and in terms of disease stage (i.e., infection duration). *Neonectria faginata* was detected in all ten sites, and *N. ditissima* in nine of ten sites, with both occurring in advancing front, killing front, and aftermath forests. Importantly, the two species regularly co-occurred, not only within the same tree (26% co-occurrence), but within the same 1-cm phloem disc (22% co-occurrence). The prevailing understanding of the dynamics BBD is that infected stands undergo a progression from initial infection by *N. ditissima* to near-total replacement by *N. faginata* in later disease stages in most cases (Houston et al., 1994; Cale et al., 2017). This idea persists despite evidence of *N. ditissima* persistence and co-occurrence of the two pathogens (Kasson and Livingston, 2009). Here we show that while *N. faginata* is the dominant species throughout killing front and aftermath forests (84% of all trees), *N. ditissima* maintained a substantial foothold throughout these stands (36% of all trees). Further, based on patterns of co-occurrence, it appears that these two species are distributed approximately randomly with respect to one another (i.e., no evidence of strong facilitation such that co-infection frequency would be elevated, nor obvious competitive exclusion within sites or trees). Previous studies have found little evidence for co-occurrence (e.g., Cale et al., 2015), but our data suggest that the two species co-occur at much higher rates than previously known.

High rates of co-occurrence were not universal, however. For example, we did find evidence that *N. faginata* becomes more prevalent relative to *N. ditissima* as disease progresses over decades. However, this appears to reflect an increase in prevalence of *N. faginata* and not the reduction or displacement of *N. ditissima*. Generally, the two species appear to have divergent climate associations, where *N. faginata* was associated with warmer climates, and *N. ditissima* with colder climates. It is possible that these patterns arise from infection duration dynamics that are obscured by our use of coarse-grained county level data. For example, northern Maine has experienced secondary killing fronts after release of beech scale populations from suppression by winter killing temperatures as temperature warms (Kasson and Livingston, 2012). However, the two warmest sites in our dataset were also of intermediate infection duration (19 and 20 years) and so infection duration is unlikely to fully explain the climate-driven distribution patterns we observed. Indeed, nongrowing season heat accumulation was the strongest predictor of *N. faginata* occurrence while infection duration was the second strongest predictor, suggesting climate as an important influence on prevalence of these species within the BBD system.

Other factors may help reconcile differences when comparing relative species prevalence from previous studies of *N. faginata* and *N. ditissima.* Stauder et al. (2020b) discussed a perithecium-dependent sampling bias, since environmental conditions and time required for perithecium production may significantly differ between these two fungi despite both being heterothallic. For example, results from a large survey of *N. faginata* and *N. ditissima* sampled from perithecia on American beech across the central Appalachian Mountains found *N. ditissima* represented just 4.2% of perithecial isolates and was recovered from only two of 13 sampled locations (Stauder et al., 2020a). This may indicate a delay in perithecium production by *N. ditissima* that favors *N. faginata* ascocarp formation or a difference in temperature optima that generally favors *N. faginata* except in the most northern stands. Further, based on our measurements of species detection rates and perithecia presence on plug periderm, if only perithecia are sampled and *N. faginata* is always detected from perithecia (assumption), then *N. ditissima* would only be detected in 1.8% of samples, whereas *N. faginata* would be detected in the remaining 98.2% of samples. This is not far from actual field detection rates. This suggests considerable bias against detecting *N. ditissima*, which is actually present in 29% of perithecial samples and 27% of all perithecial and non-perithecial samples. Another possibility is that mycoparasitic fungi detailed in this study may preferentially parasitize *N. ditissima*. Interestingly, the only site from which *Clonostachys rosea* was recovered in West Virginia in a previous study also had the highest incidence of *N. ditissima* (Stauder et al., 2020a). A second fungus, *Fusarium babinda*, may also interact differentially with *N. faginata* and *N. ditissima*. This fungus has previously been recovered from *Neonectria* perithecia (Kasson and Livingston, 2009; Stauder et al., 2020a), possibly suggesting it opportunistically colonizes these tissues (though we note that perithecia were removed from our samples prior to DNA extraction in the current study). Nevertheless, both time course studies on perithecium production and co-plating assays are needed to further resolve these relationships.

Both *Neonectria* species exhibited significant associations with various disease severity metrics. For example, both species were negatively correlated with beech scale density. Despite the fact that scale insects appear to be obligate initiating agents of BBD, Garnas et al. (2013) found negative correlations between distributions of the *Neonectria* species and scale insect. Our data appears to support this pattern. Occurrence of *N. ditissima* but not *N. faginata* was positively correlated with crown dieback class, a result supported by both HMSC modeling and indicator species analysis (ISA). This pattern resulted in increased co-infection by the two species in higher crown-dieback classes. Higher rates of co-infection associated with increasing crown dieback may indicate that a synergistic attack by the two species contributes to tree decline. For example, given different temperature associations, relatively greater activity during cooler temperatures in the nongrowing season may allow *N. ditissima* to attack hosts during periods of dormancy when defenses such as wound compartmentalization (Manion, 2003) are compromised (Copini et al., 2014). Alternatively, *N. ditissima* may be more prevalent in later stages of tree decline as a secondary pathogen that is favored by weakened host tissue (Houston, 1981; Manion, 1981). Indeed, *N. faginata* was among the top five predictors of community composition of the bark endophyte community overall, whereas *N. ditissima* was a relatively poor predictor, suggesting that colonization by *N. faginata* may be a predisposing factor for colonization of bark by a suite of other disease-associated fungi. However, the two species produced similar sized cankers on American beech in inoculation trials (Stauder et al., 2020a) suggesting that *N. ditissima* is not restricted to tissues already weakened by primary *N. faginata* infection.

### 4.3 Ecological roles of fungi in the BBD system

We used a combination of ISA and literature searches to describe the ecological roles of fungi occurring in the context of the BBD complex. ISA delineated clear groups of fungi associated with different stages of disease or disease agents, including healthy beech associates, fungi associated with intermediate or high levels of BBD pressure and tree decline, and fungi associated with presence and/or absence of the primary disease agents (i.e., *Neonectria* and beech scale). The majority of healthy beech associates were not taxonomically identified past the order level (67%). In addition, only 40% of ASVs associated with low levels of cankering and absence of beech scale were identified past the order level. Of the 24 remaining indicator ASVs 96% (23 ASVs) were taxonomically identified to at least the genus level, suggesting that bark endophytic communities of healthy American beech are a relatively unexplored reservoir of fungal diversity given comparatively low taxonomic identification. Bark endophytes have historically received less attention than foliar endophytes in temperate forests (Unterseher, 2011), and further exploration of bark endophyte diversity and functioning is warranted.

We observed a shift in both taxonomic composition and functional potential in the indicators of intermediate to high BBD pressure compared to healthy beech. Overall, 13 of the 16 ASVs (81%) associated with intermediate to high BBD pressure were annotated to saprotrophic functional groups, and five of the putatively saprotrophic ASVs associated with high BBD pressure were further identified as facultative plant pathogens. Four ASVs in intermediate- and high BBD pressure categories, including *N. faginata*, were also among the top five predictors of community composition. Many endophytes and plant pathogens function as facultative saprotrophs (Frankland, 1998; Stone et al., 2004); a shift in community function towards saprotrophy may be an important indicator of later stages of tree decline. The fungal communities associated with late stages of tree decline in particular, as indicated by high levels of crown dieback, may contribute to tree death by weakening tissues to the point of mechanical failure (Houston, 1981; Manion, 1981). Together, enrichment of saprotrophs and plant pathogens along with an apparent consistent shift in community composition indicate that the endophytic fungal community may play an important role in disease progression beyond the direct action of the primary disease agents.

We observed eight ASVs that were associated with presence or absence of the primary disease agents (*Neonectria* and beech scale). It is difficult to assign ecological roles or the nature of interactions based on pairwise species associations. For example, a positive correlation between species may indicate facilitation, overlapping habitat or microclimatic preferences, or may indicate deadlocked competition or mycoparasitic relationships (Maynard et al., 2018). Indeed, some of the ASVs in this group were associated with high beech scale density and absence of *Neonectria*, or vice versa. Two of these (ASVs 27 and 234) were restricted geographically – occurring primarily in our Wisconsin site, which, at nine years infection duration, was unique in our dataset with high average beech scale density and low *Neonectria* incidence. As such, these two ASVs may be indicators of early-stage BBD infection or geographic location rather than of interactions with beech scale or *Neonectria*, *per se*. However, two other ASVs (ASVs 69 and 217) that were geographically widespread, occurring in 4-5 sites, were both indicators of beech scale absence, with one also being an indicator of high *Neonectria* perithecium density and the other an indicator of *Neonectria* absence. It is possible that these taxa function as facultative entomopathogens/mycoparasites depending on BBD disease stage and available hosts.

We also examined distribution of two species previously reported as mycoparasites (*N. ferrugineum* and *C. rosea*) and one reported as a potential additional plant pathogen and/or an entomopathogen (*F. babinda*) in the BBD system. One of these, *N. ferrugineum* (ASV 807) occurred at only two sites and was not a statistical indicator of any disease categories. Despite its prevalence in visual surveys (Houston, 1983) our data suggest that given its relative rarity, this fungus may not play a primary role in limiting growth of *Neonectria* pathogens involved in BBD. In contrast, two ASVs were identified as *C. rosea*, which together were detected at four of our ten sites including recently infested sites in Wisconsin and Michigan, suggesting this fungus is present in BBD-affected forest stands even at the earliest stages of *Neonectria* establishment. In addition, one of the *C. rosea* ASVs was an indicator of high *Neonectria* perithecia density, consistent with its hypothesized mycoparasitic status in this system (Stauder et al., 2020a). This species has long been used as a biocontrol agent against plant pathogenic fungi (Schroers et al., 1999). Genomic mechanisms of mycoparasitism have also been described in this species (Karllson et al., 2015) making this an intriguing candidate for further study in terms of interactions with the primary *Neonectria* disease agents. *Fusarium babinda* (ASV 19) was an indicator of high BBD pressure and was found at all ten of our sites, suggesting this fungus is a geographically widespread and consistent member of late-stage decline communities associated with BBD. In particular, *F. babinda* was associated with high wax density supporting its hypothesized association with beech scale (Stauder et al., 2020a). Whether this fungus is parasitizing the scale insects remains unclear. Previous literature supports a possible entomopathogenic lifestyle, having been previously recovered from other non-native forest insect pests (*Lymantria dispar* and *Adelges tsugae*) in the eastern U.S. (Jacobs-Venter et al., 2018).

### 4.4 Significance outside the BBD pathosystem

The results of this study provide novel insights into a well-studied disease that has seen definitive, albeit incremental progress in its understanding since its discovery 130 years ago. The independent confirmation of a recently uncovered widespread fungus, *F. babinda*, associated with beech scale opens the door for functional studies and bioassays to confirm the ecology of this suspected entomopathogen. Its obscurity up until this point across BBD-impacted forests belies its relatively high incidence. Such observations highlight the importance of high-throughput amplicon sequencing in well-studied pathosystems, where causal agents are thought to be well understood.

A primary finding of this study – one that counters the conventional understanding of this system – is that *N. ditissima* is not only present in many stands but often co-occurs with *N. faginata* in the same trees, perhaps contributing to enhanced disease severity. *Neonectria faginata* is known only from beech in North America yet it has never been found outside of BBD-infected stands. Applying the findings of this study to other *Neonectria* and *Corinectria* canker disease systems has potential to uncover a native reservoir for *N. faginata*, which not unlike *N. ditissima* on beech, might be less competitive on other host substrates. Such differences in fruiting abundance are already known for *N. ditissima*: perithecium production is high on *Acer pensylvanicum* compared to limited production on *Ilex mucronata* and *Sorbus americana* and no confirmed production on *Liquidambar styraciflua* (Stauder et al., 2020a; Kasson and Stauder, unpublished observations). Since *N. ditissima* is present on over one quarter of infected beech trees, the potential for amplification of spillover of this pathogen in ways that influence non-beech hosts could also be an important mechanism by which this disease impacts forest structure, function, and diversity.

### 4.5 Conclusions

Here we described new aspects of the ecology of the two primary fungal pathogens of BBD, *N. faginata* and *N. ditissima*. In particular, *N. ditissima* occurs far more widely than previously known, co-occurring with *N. faginata* in nearly all of the sites we examined including within the same tree and even the same 1-cm phloem disc. The two species have apparently contrasting climate associations with *N. faginata* being associated with warmer temperatures and *N. ditissima* with cooler temperatures. Further, the two species appear to have different contributions to disease – our data suggest that *N. ditissima* becomes more prevalent in later stages of tree decline potentially indicating increased importance as trees progress through stages of the decline cycle. We also identified categories of fungi that may alter the trajectory of disease by functioning as entomopathogens, mycoparasites, saprotrophs and/or alternate or additional pathogens, thus causing downstream shifts in community composition in the fungal communities of beech bark.

## Acknowledgements

This work was funded in part by the New Hampshire Agricultural Experiment Station. This is Scientific Contribution Number_____. This work was supported by the USDA National Institute of Food and Agriculture McIntire-Stennis Project_____. We thank Paul Super and Bambi Teague at the National Parks Service in North Carolina as well as Keith Kanoti at the University of Maine for their help in obtaining collection permits. We also appreciate the cooperation of the New York, Michigan, Pennsylvania, and Wisconsin Departments of Natural Resources for granting collection permits.

---

**Figure S1.**
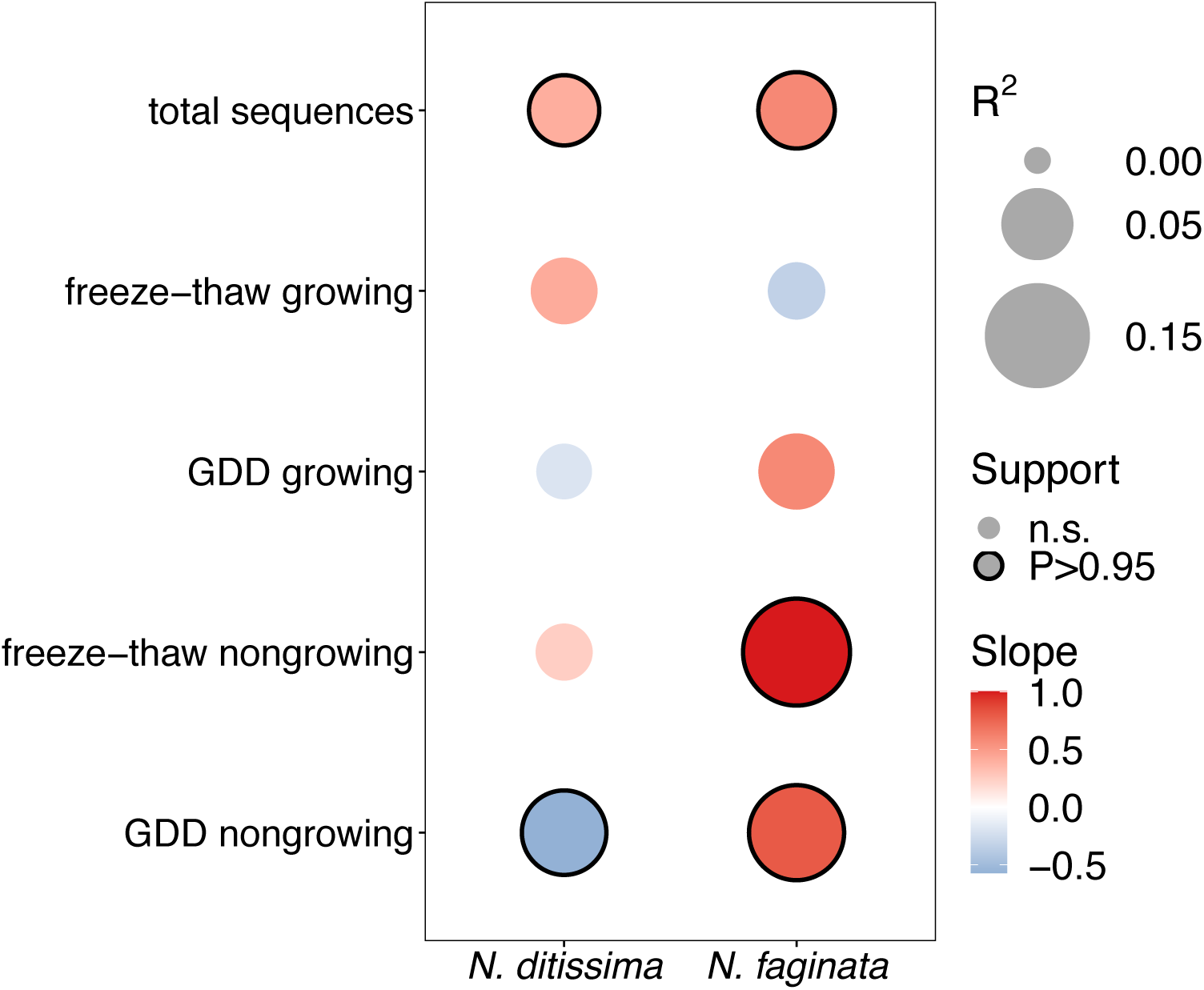
Effects of growing season and non-growing season climate on *Neonectria* distribution. Boxes with solid outlines represent effects at posterior probability *P* > 0.95 (analogous to a p-value of *P* < 0.05). Box fill indicates the strength and direction of the relationship with blue indicating negative, red indicating positive. Point size indicates the magnitude of *R*^2^.

**Figure S2.**
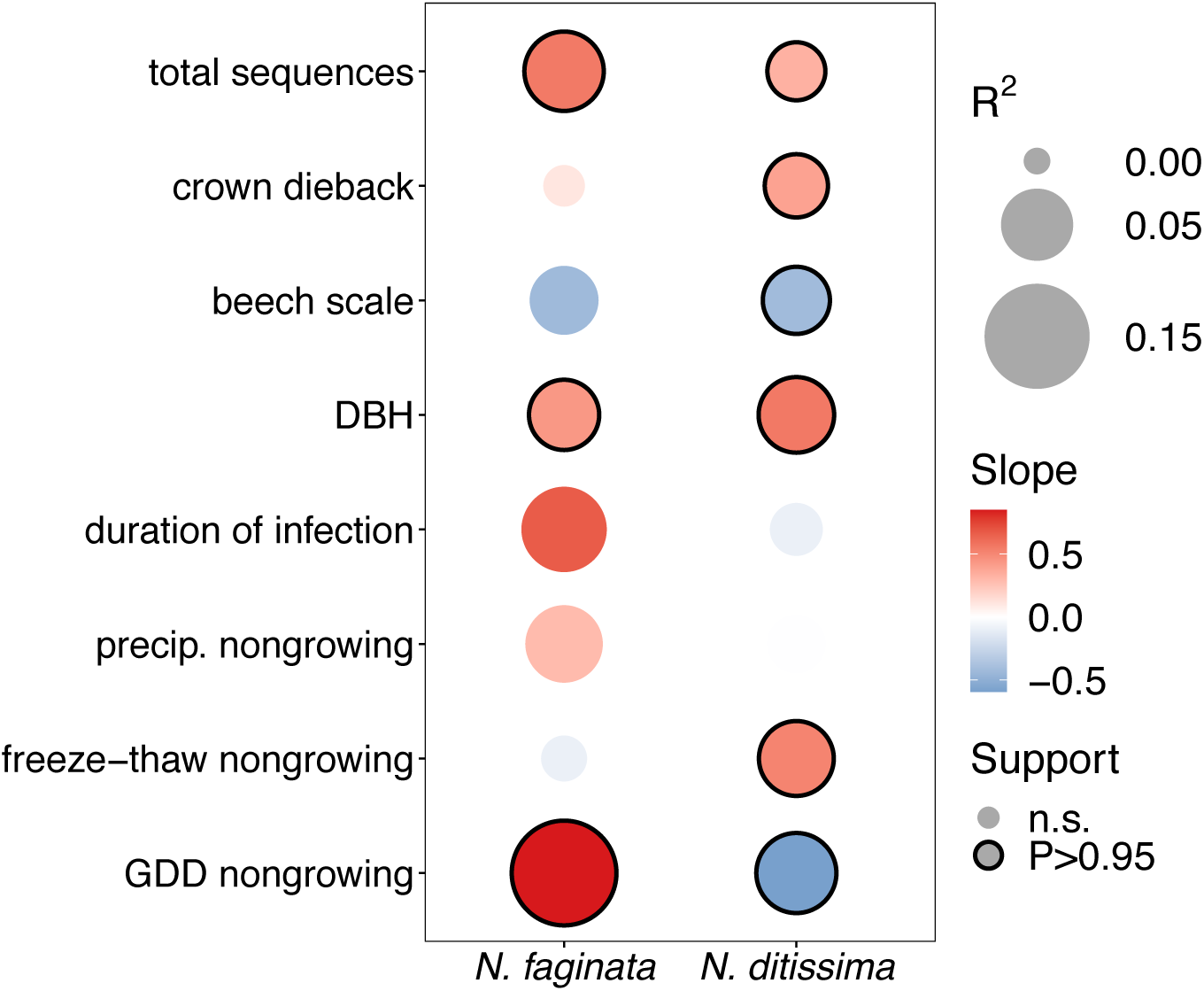
Effects of disease severity and climate on *Neonectria* species occurrence within trees. This represents a variation of the model presented in Fig. 2 with nongrowing season precipitation included in the model. Boxes with solid outlines represent significant effects at posterior probability *P* > 0.95 (analogous to a p-value of *P* < 0.05). Box fill indicates the strength and direction of the relationship with blue indicating negative, and red indicating positive. Point size indicates the magnitude of *R*^2^.

**Figure S3.**
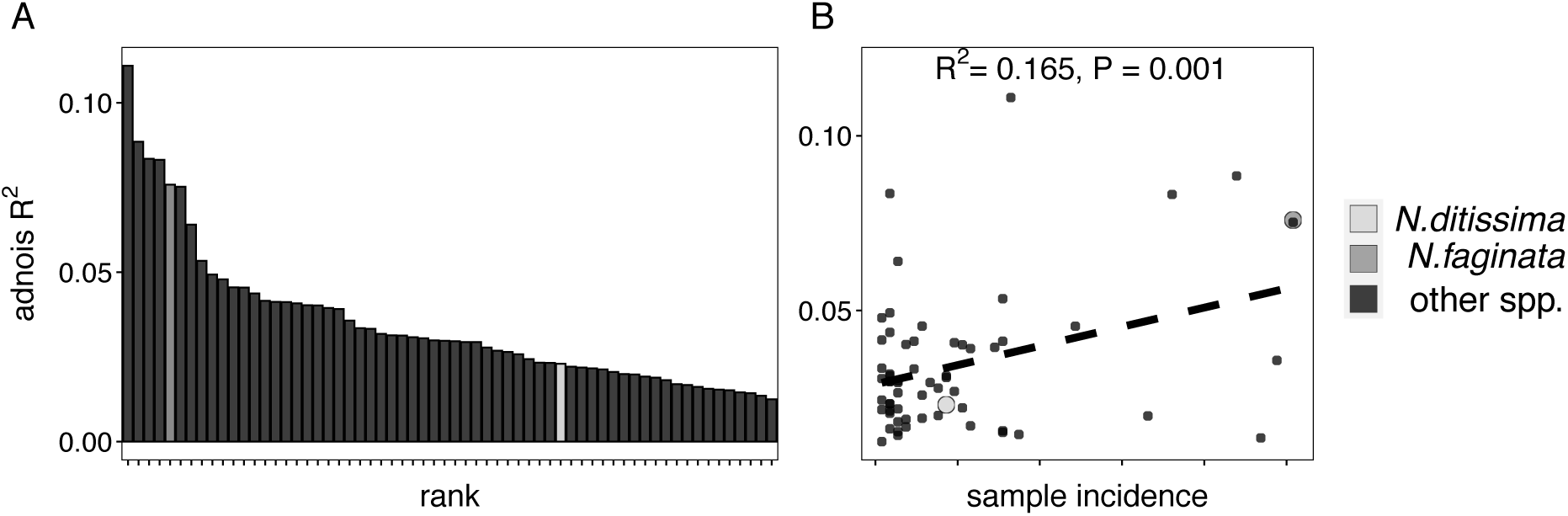
Adonis analysis of individual ASV effects on community composition (i.e., identifying which ASVs were mostly strongly associated with community turnover). Panel A: ASV frequency versus adonis *R*^2^. Panel B: Adonis *R*^2^ sorted largest to smallest. The top five ASVs in terms of adonis *R*^2^ are indicated in Table 2.

**Table S1.** Pairwise correlations of site mean disease severity, climate, and site characteristics (latitude, longitude, elevation). Correlation coefficients (Pearson’s r or Spearman’s ρ) are indicated in the lower triangle and *P* values are indicated in the upper triangle of the matrix. Correlation coefficients are based on Pearson’s r accept in the case of ordinal variables or variables with non-normal distributions (gray highlighting). Significant correlations at *P* < 0.05 are bolded.

|  | Latitude | Longitude | Elevation | GDD4<br>(growing) | GDD4<br>(nongrowing) | Freeze-thaw<br>(growing) | Freeze-thaw<br>(nongrowing) | Precipitation<br>(growing) | Precipitation<br>(nongrowing) | Infection<br>duration | DBH | Perithecia<br>density | Canker<br>density | Beech<br>scale<br>density | Tree<br>condition |
| --- | --- | --- | --- | --- | --- | --- | --- | --- | --- | --- | --- | --- | --- | --- | --- |
| Latitude |  | 0.349 | **0.016** | **0.047** | **<0.001** | 0.277 | 0.276 | **<0.001** | 0.058 | 0.273 | 0.556 | 0.385 | 0.751 | **0.033** | 0.651 |
| Longitude | 0.33 |  | **0.026** | 0.536 | 0.234 | 0.298 | 0.059 | 0.987 | 0.266 | **<0.001** | 0.128 | 0.556 | **0.003** | 0.777 | 0.907 |
| Elevation | **-0.73** | **-0.69** |  | 0.587 | **0.008** | **0.038** | 0.334 | 0.365 | 0.411 | **0.013** | 0.934 | 0.467 | 0.074 | 0.293 | 0.881 |
| GDD4 (growing) | **-0.64** | -0.22 | 0.20 |  | **0.046** | 0.968 | 0.081 | 0.328 | 0.465 | 0.602 | 0.987 | 0.150 | 0.347 | 0.803 | 0.511 |
| GDD4 (nongrowing) | **-0.95** | -0.41 | **0.78** | **0.64** |  | 0.056 | 0.450 | **0.033** | 0.076 | 0.173 | 0.881 | 0.214 | 0.425 | 0.244 | 0.244 |
| Freeze-thaw (growing) | -0.38 | -0.37 | **0.66** | 0.01 | 0.62 |  | 0.379 | 0.651 | 0.419 | 0.173 | 0.777 | 0.627 | 0.580 | 0.405 | 0.174 |
| Freeze-thaw<br>(nongrowing) | 0.38 | 0.61 | -0.34 | -0.58 | -0.27 | 0.31 |  | 0.701 | 0.516 | 0.120 | 0.425 | 0.934 | 0.108 | 0.777 | 0.627 |
| Precipitation (growing) | **-0.96** | -0.01 | 0.32 | 0.35 | **0.67** | 0.16 | -0.14 |  | 0.074 | 0.947 | 0.260 | 0.328 | 0.881 | **0.005** | 0.701 |
| Precipitation<br>(nongrowing) | -0.62 | 0.39 | 0.29 | 0.26 | 0.58 | 0.29 | 0.23 | 0.59 |  | 0.296 | **0.016** | 0.385 | 0.200 | **0.001** | 0.987 |
| Infection duration | 0.38 | **0.97** | **-0.75** | -0.19 | -0.47 | -0.47 | 0.52 | 0.02 | 0.37 |  | 0.084 | 0.300 | **0.004** | 0.638 | 0.973 |
| DBH | 0.21 | -0.52 | -0.03 | 0.01 | 0.05 | -0.10 | -0.28 | -0.39 | **-0.73** | -0.57 |  | 0.651 | 0.310 | **0.043** | 0.987 |
| Perithecia density | -0.31 | 0.21 | -0.26 | 0.49 | 0.43 | 0.18 | -0.03 | 0.35 | 0.31 | 0.36 | -0.16 |  | 0.726 | 0.533 | 0.060 |
| Canker density | 0.12 | **0.83** | -0.59 | -0.33 | -0.28 | -0.20 | 0.54 | 0.05 | 0.44 | **0.81** | -0.36 | 0.13 |  | 0.446 | 0.244 |
| Beech scale density | **0.67** | -0.10 | -0.37 | 0.09 | -0.41 | -0.30 | -0.10 | **-0.81** | **-0.88** | -0.17 | **0.65** | -0.22 | -0.27 |  | 0.934 |
| Tree condition | 0.16 | 0.04 | 0.05 | -0.24 | -0.41 | -0.47 | -0.18 | -0.14 | -0.01 | -0.01 | -0.01 | -0.61 | 0.41 | -0.03 |  |

## References

1. Barnett, H. L., and Lilly, V. G. (1962). A Destructive Mycoparasite, Gliocladium Roseum. Mycologia 54, 72–77. doi:10.1080/00275514.1962.12024980.

2. Cale, J. A., Garrison-Johnston, M. T., Teale, S. A., and Castello, J. D. (2017). Beech bark disease in North America: Over a century of research revisited. Forest Ecology and Management 394, 86–103. doi:10.1016/j.foreco.2017.03.031.

3. Cale, J. A., Teale, S. A., Johnston, M. T., Boyer, G. L., Perri, K. A., and Castello, J. D. (2015). New ecological and physiological dimensions of beech bark disease development in aftermath forests. Forest Ecology and Management 336, 99–108. doi:10.1016/j.foreco.2014.10.019.

4. Callahan, B. J., McMurdie, P. J., Rosen, M. J., Han, A. W., Johnson, A. J. A., and Holmes, S. P. (2016). DADA2: High-resolution sample inference from Illumina amplicon data. Nat Methods 13, 581–583. doi:10.1038/nmeth.3869.

5. Castlebury, L. A. C. A., Rossman, A. Y. R. Y., and Hyten, A. S. H. S. (2006). Phylogenetic relationships of Neonectria/Cylindrocarpon on Fagus in North AmericaMention of trade names or commercial products in this article is solely for the purpose of providing specific information and does not imply recommendation or endorsement by the US Department of Agriculture. Botany. doi:10.1139/b06-105.

6. Copini, P., den Ouden, J., Decuyper, M., Mohren, G. M. J., Loomans, A. J. M., and Sass-Klaassen, U. (2014). Early wound reactions of Japanese maple during winter dormancy: the effect of two contrasting temperature regimes. AoB Plants 6. doi:10.1093/aobpla/plu059.

7. Cotter H. V., and Blanchard R. O. (1981). Identification of the two *Nectria* taxa causing bole cankers on American beech. Plant Disease, 65(4):332–334.

8. Desprez-Loustau M. L., Aguayo J., Dutech C., Hayden K. J., Husson C., Jakushkin B., Marçais B., Piou D., Robin C., Vacher C. (2016). An evolutionary ecology perspective to address forest pathology challenges of today and tomorrow. Annals of Forest Science. 73(1):45–67.

9. De Caceres, M., Legendre, P. (2009). Associations between species and groups of sites: indices and statistical inference. Ecology, doi: 10.1890/08-1823.1

10. Dukes, J. S. D. S., Pontius, J. P., Orwig, D. O., Garnas, J. R. G. R., Rodgers, V. L. R. L., Brazee, N. B., et al., (2009). Responses of insect pests, pathogens, and invasive plant species to climate change in the forests of northeastern North America: What can we predict?This article is one of a selection of papers from NE Forests 2100: A Synthesis of Climate Change Impacts on Forests of the Northeastern US and Eastern Canada. Canadian Journal of Forest Research. doi:10.1139/X08-171.

11. Edgar, R. C. (2013). UPARSE: highly accurate OTU sequences from microbial amplicon reads. Nat Methods 10, 996–998. doi:10.1038/nmeth.2604.

12. Ehrlich, J. (2011). The beech bark disease: a *Nectria* disease of *Fagus*, following *Cryptococcus fagi* (BAER.). Canadian Journal of Research. doi:10.1139/cjr34-070.

13. Feau, N., and Hamelin, R. C. (2017). Say hello to my little friends: how microbiota can modulate tree health. New Phytologist 215, 508–510. doi:https://doi.org/10.1111/nph.14649.

14. Frankland, J. C. (1998). Fungal succession — unravelling the unpredictable. Mycological Research 102, 1–15. doi:10.1017/S0953756297005364.

15. Frøslev, T. G., Kjøller, R., Bruun, H. H., Ejrnæs, R., Brunbjerg, A. K., Pietroni, C., et al., (2017). Algorithm for post-clustering curation of DNA amplicon data yields reliable biodiversity estimates. Nat Commun 8, 1–11. doi:10.1038/s41467-017-01312-x.

16. Garnas, J. R., Ayres, M. P., Liebhold, A. M., and Evans, C. (2011b). Subcontinental impacts of an invasive tree disease on forest structure and dynamics. Journal of Ecology 99, 532–541. doi:https://doi.org/10.1111/j.1365-2745.2010.01791.x.

17. Garnas, J. R., Houston, D. R., Ayres, M. P., and Evans, C. (2011a). Disease ontogeny overshadows effects of climate and species interactions on population dynamics in a nonnative forest disease complex. Ecography 35, 412–421. doi:https://doi.org/10.1111/j.1600-0587.2011.06938.x.

18. Garnas, J. R., Houston, D. R., Twery, M. J., Ayres, M. P., and Evans, C. (2013). Inferring controls on the epidemiology of beech bark disease from spatial patterning of disease organisms. Agricultural and Forest Entomology 15, 146–156. doi:https://doi.org/10.1111/j.1461-9563.2012.00595.x.

19. Gómez-Cortecero, A., Saville, R. J., Scheper, R. W. A., Bowen, J. K., Agripino De Medeiros, H., Kingsnorth, J., et al., (2016). Variation in Host and Pathogen in the Neonectria/Malus Interaction; toward an Understanding of the Genetic Basis of Resistance to European Canker. Front Plant Sci 7. doi:10.3389/fpls.2016.01365.

20. Griffith, D. M., Veech, J. A., and Marsh, C. J. (2016). cooccur: Probabilistic Species Co-Occurrence Analysis in R. Journal of Statistical Software 69, 1–17. doi:10.18637/jss.v069.c02.

21. Harrell, F.E. Jr, Dupont, C. et al., (2019). Hmisc: Harrell Miscellaneous. R package version 4.3-0. https://CRAN.R-project.org/package=Hmisc

22. Houston, D. R. (1981). Stress Triggered Tree Diseases: The Diebacks and Declines. U.S. Department of Agriculture, Forest Service.

23. Houston, D. R. (1983). Effects of parasitism by Nematogonum ferrugineum (Gonstorrhodiella highlei) on pathogenicity of Nectria coccinea var. faginata and Nectria galligena. In: Proceedings, I.U.F.R.O. Beech Bark Disease Working Party Conference; 1982 September 26-October 8; Hamden, CT. Sponsored by the USDA Forest Service, Northeastern Forest Experiment Station. Gen. Tech. Rep. WO-37. [Washington, DC]: U.S. Department of Agriculture, Forest Service: 109-114. 37, 109–114.

24. Houston, D.R., Mahoney, E.M. and McGauley, B.H., 1987, November. Beech bark disease – association of *Nectria ochroleuca* in W VA, PA, and Ontario. In Phytopathology (Vol. 77, No. 11, pp. 1615–1615). 3340 Pilot Knob Road, St. Paul, MN 55121: Amer Phytopathological Soc.

25. Houston, D. R. (1994). Major new tree disease epidemics: beech bark disease. Annu. Rev. Phytopathol. 32, 75–87. doi:10.1146/annurev.py.32.090194.000451.

26. Houston, D. R., Rubin, B. D., Twery, M. J., Steinman, J. R., and Steinman, J. R. (2005). Spatial and Temporal Development of Beech Bark Disease in the Northeastern United States. In: Evans, Celia A., Lucas, Jennifer A. and Twery, Mark J., eds. Beech Bark Disease: Proceedings of the Beech Bark Disease Symposium; 2004 June 16-18; Saranak Lake, NY. Gen. Tech. Rep. NE-331. Newtown Square, PA: US. Department of Agriculture, Forest Service, Northeastern Research Station: 43–47. Available at: https://www.fs.usda.gov/treesearch/pubs/20405 [Accessed November 24, 2020].

27. Houston, D. R., and Valentine, H. T. (2011). Beech bark disease: the temporal pattern of cankering in aftermath forests of Maine. Canadian Journal of Forest Research. doi:10.1139/x88-007.

28. Jacobs-Venter, A., Laraba, I., Geiser, D. M., Busman, M., Vaughan, M. M., Proctor, R. H., et al., (2018). Molecular systematics of two sister clades, the *Fusarium concolor* and *F. babinda* species complexes, and the discovery of a novel microcycle macroconidium–producing species from South Africa. Mycologia 110, 1189–1204. doi:10.1080/00275514.2018.1526619.

29. Karlsson, M., Durling, M. B., Choi, J., Kosawang, C., Lackner, G., Tzelepis, G. D., et al., (2015). Insights on the Evolution of Mycoparasitism from the Genome of *Clonostachys rosea*. Genome Biol Evol 7, 465–480. doi:10.1093/gbe/evu292.

30. Kasson, M. T., and Livingston, W. H. (2009). Spatial distribution of *Neonectria* species associated with beech bark disease in northern Maine. Mycologia 101, 190–195. doi:10.3852/08-165.

31. Kasson, M. T., and Livingston, W. H. (2012). Relationships among beech bark disease, climate, radial growth response and mortality of American beech in northern Maine, USA. Forest Pathology 42, 199–212. doi:https://doi.org/10.1111/j.1439-0329.2011.00742.x.

32. Kolp, M., Double, M. L., Fulbright, D. W., MacDonald, W. L., and Jarosz, A. M. (2020). Spatial and temporal dynamics of the fungal community of chestnut blight cankers on American chestnut (Castanea dentata) in Michigan and Wisconsin. Fungal Ecology 45, 100925. doi:10.1016/j.funeco.2020.100925.

33. Manion, P. D. (1981). Tree disease concepts. Tree disease concepts. Available at: https://www.cabdirect.org/cabdirect/abstract/19810672031 [Accessed February 10, 2021].

34. Manion, P. D. (2003). Evolution of Concepts in Forest Pathology. Phytopathology® 93, 1052–1055. doi:10.1094/PHYTO.2003.93.8.1052.

35. Maynard, D. S., Covey, K. R., Crowther, T. W., Sokol, N. W., Morrison, E. W., Frey, S. D., et al., (2018). Species associations overwhelm abiotic conditions to dictate the structure and function of wood-decay fungal communities. Ecology 99, 801–811. doi:10.1002/ecy.2165.

36. Morin, R. S. M. S., Liebhold, A. M. L. M., Tobin, P. C. T. C., Gottschalk, K. W. G. W., and Luzader, E. L. (2007). Spread of beech bark disease in the eastern United States and its relationship to regional forest composition. Canadian Journal of Forest Research. doi:10.1139/X06-281.

37. Nilsson, R. H., Larsson, K.-H., Taylor, A. F. S., Bengtsson-Palme, J., Jeppesen, T. S., Schigel, D., et al., (2019). The UNITE database for molecular identification of fungi: handling dark taxa and parallel taxonomic classifications. Nucleic Acids Res 47, D259–D264. doi:10.1093/nar/gky1022.

38. Oksanen, J., F. Blanchet, G., Friendly, M., Kindt, R., Legendre, P., McGlinn, D., et al., (2019). vegan: Community Ecology Package. R package version 2.5–6. https://CRAN.R-project.org/package=vegan

39. Ovaskainen, O., Roy, D. B., Fox, R., and Anderson, B. J. (2016). Uncovering hidden spatial structure in species communities with spatially explicit joint species distribution models. Methods in Ecology and Evolution 7, 428–436. doi:https://doi.org/10.1111/2041-210X.12502.

40. Ovaskainen, O., Tikhonov, G., Norberg, A., Blanchet, F. G., Duan, L., Dunson, D., et al., (2017). How to make more out of community data? A conceptual framework and its implementation as models and software. Ecology Letters 20, 561–576. doi:https://doi.org/10.1111/ele.12757.

41. Pauvert, C., Buée, M., Laval, V., Edel-Hermann, V., Fauchery, L., Gautier, A., et al., (2019). Bioinformatics matters: The accuracy of plant and soil fungal community data is highly dependent on the metabarcoding pipeline. Fungal Ecology 41, 23–33. doi:10.1016/j.funeco.2019.03.005.

42. PRISM Climate Group, Oregon State University, http://prism.oregonstate.edu, created May 28, 2020

43. Põlme, S., Abarenkov, K., Nilsson, H. R., Lindahl, B. D., Clemmensen, K. E., Kauserud, H., et al., (2021). FungalTraits: a user-friendly traits database of fungi and fungus-like stramenopiles. Fungal Diversity. doi:10.1007/s13225-020-00466-2.

44. Richardson, A. D., Bailey, A. S., Denny, E. G., Martin, C. W., and O’keefe, J. (2006). Phenology of a northern hardwood forest canopy. Global Change Biology 12, 1174–1188. doi:10.1111/j.1365-2486.2006.01164.x.

45. Rivers, A. R., Weber, K. C., Gardner, T. G., Liu, S., and Armstrong, S. D. (2018). ITSxpress: Software to rapidly trim internally transcribed spacer sequences with quality scores for marker gene analysis. F1000Res 7, 1418. doi:10.12688/f1000research.15704.1.

46. Rognes, T., Flouri, T., Nichols, B., Quince, C., and Mahé, F. (2016). VSEARCH: a versatile open source tool for metagenomics. PeerJ 4, e2584. doi:10.7717/peerj.2584.

47. Rossman, A. Y., Seifert, K. A., Samuels, G. J., Minnis, A. M., Schroers, H.-J., Lombard, L., et al., (2013). Genera in Bionectriaceae, Hypocreaceae, and Nectriaceae (Hypocreales) proposed for acceptance or rejection. IMA Fungus 4, 41–51. doi:10.5598/imafungus.2013.04.01.05.

48. Schroers, H.-J., Samuels, G. J., Seifert, K. A., and Gams, W. (1999). Classification of the mycoparasite Gliocladium roseum in Clonostachys as C. rosea, its relationship to Bionectria ochroleuca, and notes on other Gliocladium-like fungi. Mycologia 91, 365–385. doi:10.1080/00275514.1999.12061028.

49. Shapiro, S. S., and Wilk, M. B. (1965). An Analysis of Variance Test for Normality (Complete Samples). Biometrika 52, 591–611. doi:10.2307/2333709.

50. Shigo, A. L. (1972). The Beech Bark Disease Today in the Northeastern U.S. Journal of Forestry 70, 286–289. doi:10.1093/jof/70.5.286.

51. Stauder, C. M., Garnas, J. R., Morrison, E. W., Salgado-Salazar, C., and Kasson, M. T. (2020b). Characterization of mating type genes in heterothallic Neonectria species, with emphasis on N. coccinea, N. ditissima, and N. faginata. Mycologia 112, 880–894. doi:10.1080/00275514.2020.1797371.

52. Stauder, C. M., Utano, N. M., and Kasson, M. T. (2020a). Resolving host and species boundaries for perithecia-producing nectriaceous fungi across the central Appalachian Mountains. Fungal Ecology 47, 100980. doi:10.1016/j.funeco.2020.100980.

53. Stone, J. K., Polishook, J. D., and White, J. F. (2004). “Endophytic fungi,” in Biodiversity of Fungi, eds. M. Foster and G. Bills (Burlington: Elsevier Academic Press), 241–270. doi:10.13140/RG.2.1.2497.0726.

54. Taylor, D. L., Walters, W. A., Lennon, N. J., Bochicchio, J., Krohn, A., Caporaso, J. G., et al., (2016). Accurate Estimation of Fungal Diversity and Abundance through Improved Lineage-Specific Primers Optimized for Illumina Amplicon Sequencing. Appl. Environ. Microbiol. 82, 7217–7226. doi:10.1128/AEM.02576-16.

55. Tikhonov, G., Abrego, N., Dunson, D., and Ovaskainen, O. (2017). Using joint species distribution models for evaluating how species-to-species associations depend on the environmental context. Methods in Ecology and Evolution 8, 443–452. doi:10.1111/2041-210X.12723.

56. Tjur, T. (2009). Coefficients of Determination in Logistic Regression Models—A New Proposal: The Coefficient of Discrimination. The American Statistician 63, 366–372. doi:10.1198/tast.2009.08210.

57. Unterseher, M. (2011). “Diversity of Fungal Endophytes in Temperate Forest Trees,” in Endophytes of Forest Trees: Biology and Applications Forestry Sciences., eds. A. M. Pirttilä and A. C. Frank (Dordrecht: Springer Netherlands), 31–46. doi:10.1007/978-94-007-1599-8_2.

58. Veech, J. A. (2013). A probabilistic model for analysing species co-occurrence. Global Ecology and Biogeography 22, 252–260. doi:https://doi.org/10.1111/j.1466-8238.2012.00789.x.

59. Wingfield, M. J., Garnas, J. R., Hajek, A., Hurley, B. P., de Beer, Z. W., and Taerum, S. J. (2016). Novel and co-evolved associations between insects and microorganisms as drivers of forest pestilence. Biol Invasions 18, 1045–1056. doi:10.1007/s10530-016-1084-7.

60. Wingfield, M. J., Slippers, B., and Wingfield, B. D. (2010). Novel associations between pathogens, insects and tree species threaten world forests. New Zealand Journal of Forestry Science, 9.

